# SV^2^: Accurate Structural Variation Genotyping and *De Novo* Mutation Detection from Whole Genomes

**DOI:** 10.1101/113498

**Authors:** Danny Antaki, William M Brandler, Jonathan Sebat

## Abstract

**Motivation:** Structural Variation (SV) detection from short-read whole genome sequencing is error prone, presenting significant challenges for population or family-based studies of disease.

**Results:** Here we describe SV^2^, a machine-learning algorithm for genotyping deletions and duplications from paired-end sequencing data. SV^2^ can rapidly integrate variant calls from multiple structural variant discovery algorithms into a unified call set with high genotyping accuracy and capability to detect de novo mutations.

**Availability and Implementation:** SV^2^ is freely available on GitHub (https://github.com/dantaki/SV2)

## 1 Introduction

Structural variation (SV) is defined as a change of the structure of a chromosome larger than 50bp. SV is a major contributor to human genetic variation and is also implicated in a variety of human diseases (Conrad, et al., 2010; Sudmant, et al., 2015). In particular, *de novo* structural mutations (those in offspring and not in parents) contribute significant risk for idiopathic autism and intellectual disability (Brandler, et al., 2016). Accurate detection of *de novo* SVs is challenging due to genotyping errors, such as false-positives in offspring and false-negatives in parents. Additionally, genotyping errors bias the transmission of variants from parent to offspring, confounding analysis of pedigrees.

Characterizing SV from next generation sequencing is a difficult task due to the broad range of SV sizes (50 bp-50 Mb) and types. Hence, to fully capture the diversity of SVs a multitude of tools is required, each designed as a standalone solution (GoNL Consortium, 2014; Sudmant, et al., 2015). Methods for harmonizing variant calls and confidence scores from multiple SV calling methods into a unified set of SV genotypes are lacking.

## 2 Input and Methods

SV^2^ (support-vector structural-variant genotyper) is an open source application written in Python that requires a BAM file, a single nucleotide variant (SNV) VCF file, and either a BED or VCF file of deletions and duplications as input. SV^2^ operates in three stages: preprocessing, feature extraction, followed by genotyping (Fig. 1). Genotyping utilizes four informative features: depth of coverage, discordant paired-ends, split-reads, and heterozygous allele ratio. Genotyping is performed with supervised support vector machine classifiers trained on whole genome sequences (WGS) from the 1000 Genomes Project (1KGP). The resulting VCF file of genotypes contains annotations for genes, repeat elements, and variant identifiers of common SVs recorded by the 1KGP (Sudmant, et al., 2015).

**Figure 1:**
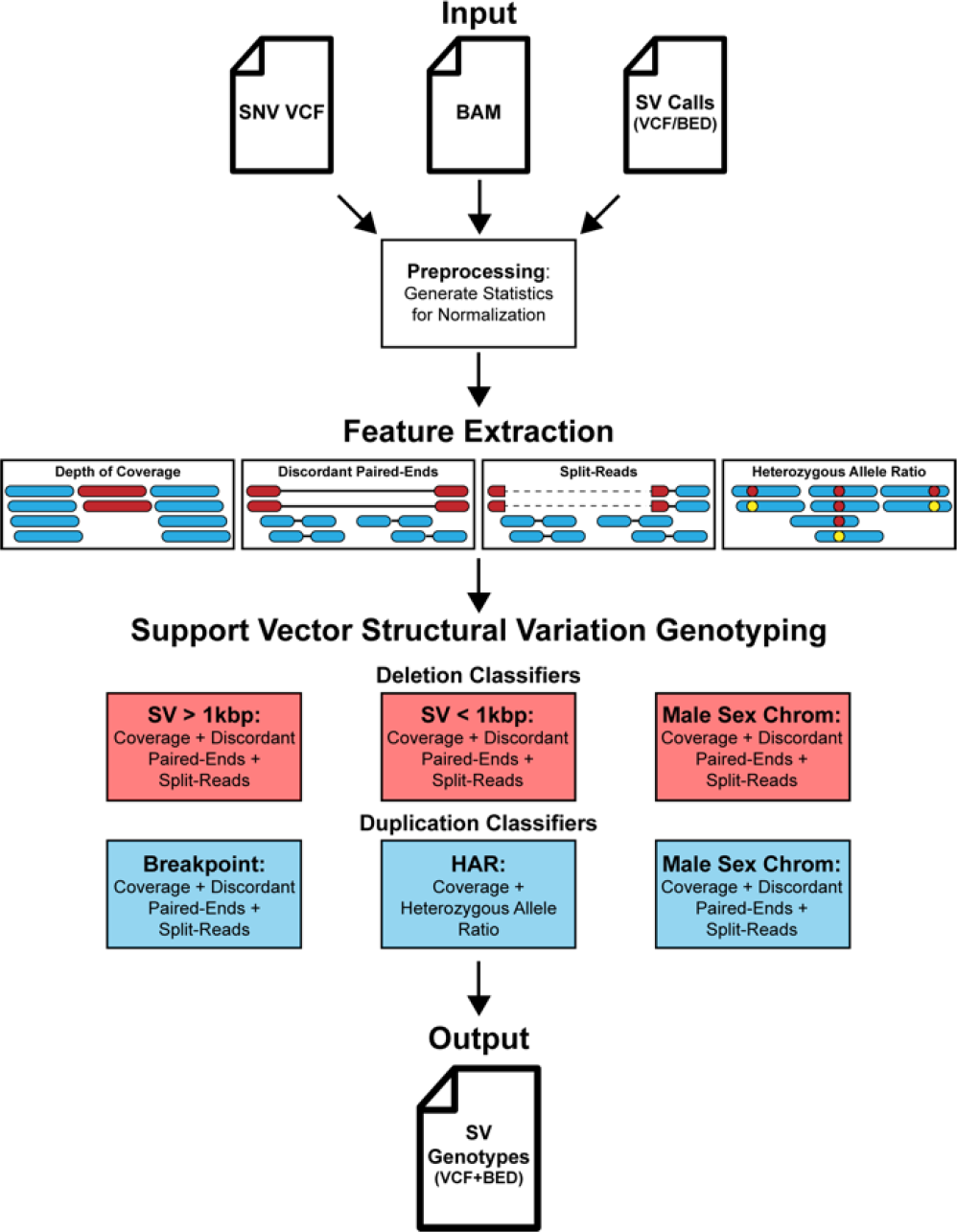
SV^2^ workflow. SV^2^ requires a VCF file of SNVs, a BAM file, and a set of SVs to genotype as input. Before genotyping, preprocessing is performed where the median coverage, insert size, and read length is recorded for feature normalization. Features for genotyping, which include depth of coverage, discordant paired-ends, split-reads, and heterozygous allele ratio (HAR), are measured for each SV. SVs are then genotyped with an ensemble of support vector machine classifiers. SV^2^ produces two output files, a BED file and a VCF, containing annotations for RefSeq genic elements, Repeat-Masker repeats, Segmental Duplications, Short Tandem Repeats, and common SVs from the 1000 Genomes phase 3 call set.

## 3 Genotyping Classifiers

Our training set included high coverage (48x, N=27) and low coverage (7x, N=2,494) WGS from the 1KGP. Training features were collected from a gold-standard SV call set on the above individuals with an estimated false discovery rate (FDR) of 1-4% (Sudmant, et al., 2015), totaling 297,131 genotypes from 11,747 unique loci. (Supplementary Table. S1).

An ensemble of SV detection methods is required to capture a wide diversity of SVs, resulting in a set of SVs with variable characteristics. Deletions have a very broad size range (50 bp->10Mb), and as deletions become very small, the signal from discordant paired-ends diminishes and signal from split-reads increases. Duplication calls, by contrast, are larger on average (>3kb Sudmant, et al., 2015) with less precise breakpoints. In addition, genotyping methods must account for the ploidy of sex chromosomes in males and females. Consequently, a one-size-fits-all genotyping method performs poorly on the combined calls

With our goal of combining SV calls from multiple methods and generating a uniform set of genotypes, we developed a genotyper consisting of multiple classifiers, each designed to genotype a different category of SV. To contend with the variable characteristics described above, deletions were stratified by size and duplications by the availability of breakpoint features (discordant paired-ends or split-reads). Deletions were trained in three classifiers: (1) deletions greater than 1kb, (2) deletions less than or equal to 1kb, and (3) deletions on male sex chromosomes. All three deletion classifiers implemented coverage, discordant paired-end, and split-read features extracted from high coverage WGS. Duplications were trained in three additional classifiers: (4) duplications with the breakpoint features, (5) duplications without breakpoint features, (6) duplications on male sex chromosomes. Classifier #5 was unique in that it included heterozygous SNV allele ratios (HAR) as a feature (“HAR” Fig. 1). The deletion male sex chromosomes classifier was not split by size due to the sparse number of training examples (N=191, Supplementary Table S1).

## 4 Performance Results

For this study, we implemented three independent WGS datasets for training, evaluation and validation (Supplementary Table S2). Parameter selection for classifiers was based on a 7-fold cross-validation of the training set (Supplementary Fig. S1) and further based on classifier performance in an independent dataset consisting of 42x WGS of 57 subjects (Supplementary Table S3), a subset of samples from our previous study of autism (“evaluation dataset”, Brandler, et al., 2017). Genotyping accuracy was evaluated further based on rates of SV transmission in families from an additional 1,827 genomes from Brandler et al. Finally, classifier performance was independently evaluated in a third dataset consisting of 72x Illumina WGS of 9 subjects from the 1KGP (“validation dataset”). We confirmed SV^2^ genotypes in the evaluation and validation datasets using two orthogonal platforms: Illumina 2.5M microarrays and Pacific Biosciences (PacBio) single molecule sequencing, respectively.

Performance was evaluated at two levels of SV filtering stringency: a “standard” level and a stricter level for calling *de novo* mutations, which are enriched for false-positive calls. Our formulation of filters applied the evaluation set of 57 subjects with SV^2^ genotypes from SV calls from ForestSV (v0.3.3 Michaelson and Sebat, 2012), LUMPY (v0.2.13 Layer, et al., 2014), and Manta (v0.2.13 Layer, et al., 2014). We implemented Illumina 2.5M arrays to calculate the false discovery rate (FDR) using SVToolkit (v2.0 sourceforge.net/projects/svtoolkit), which performs an *in silico* validation by ranking microarray probes intensities, defining the FDR as two times the fraction of variants with p-value >0.5 (Sudmant, et al., 2015). The final thresholds for filters for each stringency level considered variant length, feature availability, and FDR (Supplementary Table S4).

### Evaluation

Using the evaluation set mentioned above, we found a FDR of 40% for both unfiltered deletions (N=5,344) and duplications (N=776) (Fig. 2A). Filtering at the standard level of stringency reduced the FDR to 1.24% for deletions and 4.41% for duplications (Supplementary Fig. S2). We then ascertained the FDR of unfiltered *de novo* variants to be 60% for deletions and 86% for duplications. Applying the *de novo* filters reduced the FDR to 0.54% for deletions and 0% for duplications (Supplementary Fig. S2). We extended our evaluation of SV^2^ genotyping with an additional 1,827 individuals to the previous evaluation set of 57 subjects, totaling 619 families (N=1,884). Validation of SV^2^ genotypes implemented the group-wise transmission disequilibrium test (Chen, et al., 2015), which tests for deviations from the expected variant transmission rate (50%). An under-transmission of variants indicates either an enrichment of either false positives in parents or false negatives in the offspring. As expected, unfiltered SV calls exhibited a significant under-transmission bias: transmission rates of 39.8% for deletions and 35.08% for duplications (deletions: P=9.61x10-51, N=105,023; duplications P=7.8x10-18, N=346,173) (Fig. 2B). Applying standard genotype likelihood filters reduced the transmission bias to 48.2% (P=1.32x10-2, N=40,587) for deletions and 47.3% (P=3.39x10-3, N=3,863) for duplications. Applying more stringent *de novo* filters further reduced under-transmission bias to 49.1% (P=1.32x10-2, N=21,772) for deletions and 49.3% (P=1.0, N=2,847) for duplications (Supplementary Fig. S3).

**Figure 2:**
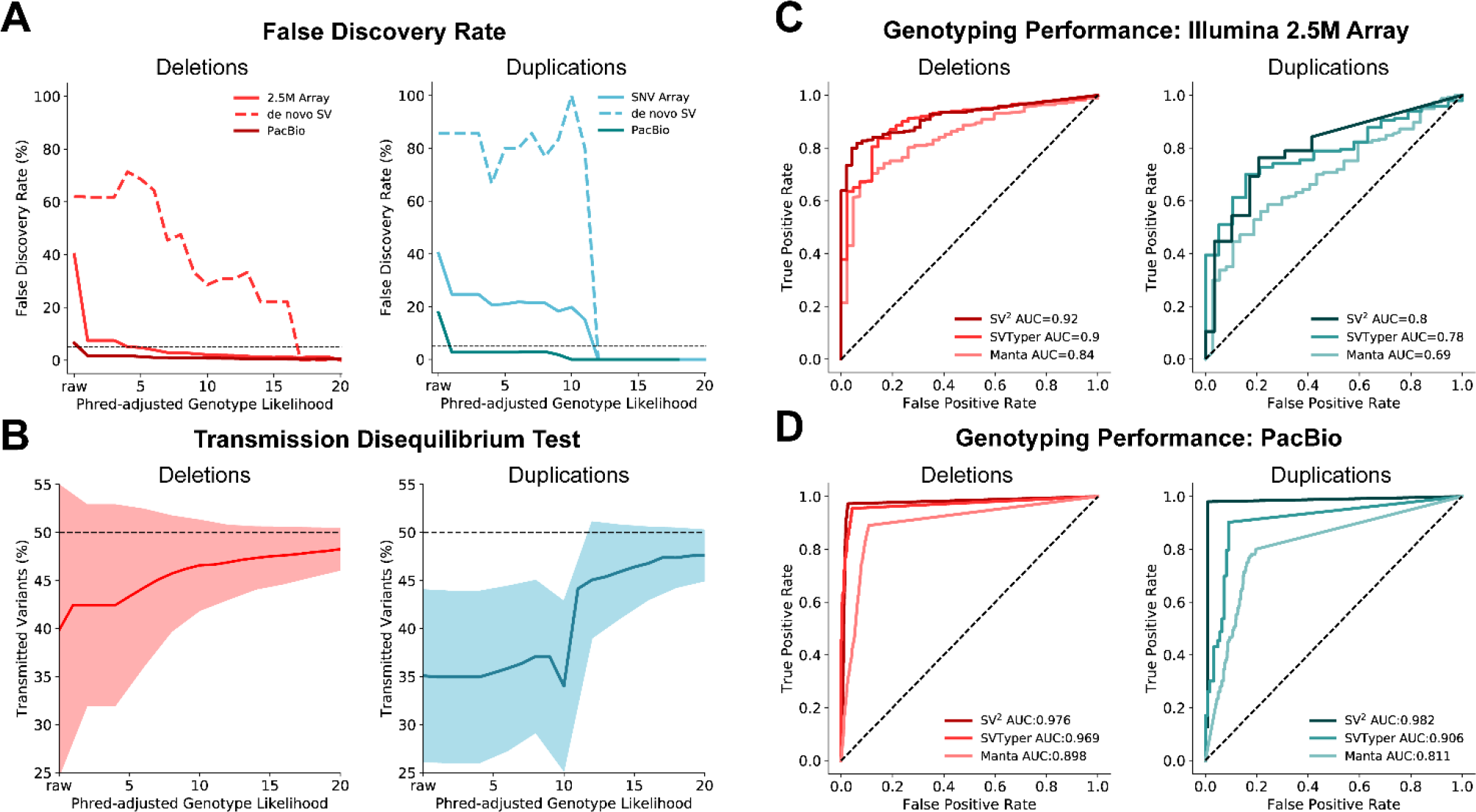
SV^2^ genotyping performance. **A**: False discovery rate across SV^2^ genotype likelihoods estimated from Illumina 2.5M arrays (N=57) and PacBio long reads (N=9). Black dotted line indicates 5% FDR. **B**: Group-wise transmission disequilibrium tests across SV^2^ genotype likelihoods in 630 offspring with shaded regions representing one standard deviation. **C**: Receiver operating characteristic (ROC) curves of WGS genotyping calculated from Illumina 2.5M arrays for SV^2^, SVTyper, and Manta in 57 individuals. **D:** ROC curves of WGS genotyping calculated from supporting PacBio SMRT reads for SV^2^, SVTyper, and Manta for SVs in 9 individuals.

Performance of SV^2^ was then compared to that of two widely used SV genotyping software SVTyper (v0.0.4 Chiang, et al., 2015) and Manta. For this comparison, SVTyper genotyped SV predictions using the companion tool LUMPY, Manta produced genotypes for its predictions, and SV^2^ genotyped the union of LUMPY and Manta calls for the previous evaluation set of 57 subjects with Illumina 2.5M arrays. Receiver operating characteristic (ROC) curves for each genotyping method were generated, specifying true and false positives with SVToolkit. SV^2^ achieved the best genotyping accuracy with an AUC of 0.92 for deletions and 0.8 for duplications, in contrast to Manta (deletion AUC=0.84, duplication AUC=0.69) and SVTyper (deletion AUC=0.9, duplications AUC=0.78) (Fig. 2C).

### Validation

Further assessment of SV^2^’s genotype likelihood filters leveraged PacBio long-read WGS (x26) on 9 subjects from the 1KGP. This validation set is independent from the training set since SV^2^ genotypes were generated using a separate deep (x72) Illumina WGS library with SV predictions from LUMPY and Manta, both of which were not implemented in the training call set (Sudmant, et al., 2015). To comply with the data release requirements for these data, only variants on chromosome 1 were analyzed. As a precaution for overfitting, we excluded SVs that overlapped with >=80% reciprocal overlap to SVs in our training set. Additionally, we omitted variants with less than 3 PacBio reads within 1kbp flanking regions. Valid WGS genotypes required at least one supporting breakpoint with 50% reciprocal overlap to a PacBio split-read or CIGAR string. The FDR was 6.53% (N=3,121) and 17.72% (N=413) for unfiltered deletions and duplications respectively (Fig. 2A). SV^2^ standard filters, lowered the FDR for deletions to 0.85% (*de novo* filters: 0.62%) and for duplications to 0% (*de novo* filters: 0%). With these data, we then compared SV^2^ genotyping performance to the aforementioned genotyping methods. Likewise, we found that SV^2^ produced the optimal performance with AUCs of 0.98 for deletions and duplications. Conversely, Manta performance resulted in an AUC of 0.9 for deletions and 0.81 for duplications, and SVTyper producing AUCs of 0.97 for deletions and 0.91 for duplications (Fig. 2D).

## 5 Conclusions

SV^2^ is unique from other genotyping methods in its use of machine learning, specifically a radial basis function kernel which is able to distinguish classes among nonlinear distributions (Supplementary Fig. S4). SV^2^ can rapidly genotype a wide variety of deletions and duplications (Supplementary Fig. S5). Exceptions include variants that completely overlap segmental duplications, short tandem repeats, centromeres, telomeres, or other unsequenceable regions due to complications with unique-mappability from short-read technology. SV^2^ is designed for genotyping SVs in population genetic studies, including studies of complex traits or disease in pedigrees or case-control samples. SV^2^ could be further applied to any comparison of SVs across samples, for example in identifying somatic variants from multiple genomes derived from individual cells or clones of one individual (Abyzov, et al., 2012). SV^2^ aids in variant post-processing by recording overlap to common filtering criteria and gene elements, and provides the option to merge divergent breakpoints according to the optimal genotype likelihood. Ultimately, SV^2^’s strength is its ability to harmonize SV predictions from multiple callers, simplifying genotyping, likelihood estimation, analysis of SV association, and providing a much-needed tool for accurately detecting *de novo* mutations.

## Acknowledgements

We would like to thank the 1000 Genomes Project and the Human Genome Structural Variation Consortium for providing PacBio SMRT alignments and complementary high coverage PCR-free Illumina short-read alignments for chromosome 1.

